# Chromosome level genome reference of the Caucasian dwarf goby *Knipowitschia* cf. *caucasica*, a new alien Gobiidae invading the River Rhine

**DOI:** 10.1101/2024.04.22.590508

**Authors:** Alexandra Schoenle, Nadège Guiglielmoni, Tobias Mainz, Carola Greve, Alexander Ben Hamadou, Lisa Heermann, Jost Borcherding, Ann-Marie Waldvogel

## Abstract

The Caucasian dwarf goby *Knipowitschia* cf. *caucasica* is a new invasive alien Gobiidae spreading in the Lower Rhine since 2019. Little is known about the invasion biology of the species and further investigations to reconstruct the invasion history are lacking genomic resources. We assembled a high-quality chromosome-scale reference genome of *Knipowitschia* cf. *caucasica* by combining PacBio, Omni-C and Illumina technologies. The size of the assembled genome is 956.58 Mb with a N50 scaffold length of 43 Mb, which includes 92.3 % complete Actinopterygii Benchmarking Universal Single-Copy Orthologs. 98.96 % of the assembly sequence was assigned to 23 chromosome-level scaffolds, with a GC-content of 42.83 %. Repetitive elements account for 53.08 % of the genome. The chromosome-level genome contained 26,404 transcripts with 23,210 multi-exons, of which 26,260 genes were functionally annotated. In summary, the high-quality genome assembly provides a fundamental basis to understand the adaptive advantage of the species.

## Introduction

Starting in 1999, few years after the opening of the Rhine-Main-Danube-Channel, a continuous succession of four gobiid fish invasions has been documented in the River Rhine, particularly the Lower Reaches (Borcherding, Staas, et al., 2011). The fifth and most recent invasion of the Caucasian dwarf goby *Knipowitschia* cf. *caucasica* is exceeding the preceeding goby invasions both in the rate of population growth and in competition with the resident fish community including native and invasive species (Borcherding, Aschemeier, et al., 2021, and unpublished data from catches in 2022 by Lisa Heermann and Jost Borcherding). *Knipowitschia* cf. *caucasica* ranks to the highest trophic level in the ecological food chain of the aquatic habitat, feeding on zooplankton and small chironomid larvae from macrozoobenthos (Borcherding, Aschemeier, et al., 2021; Didenko et al., 2020). It is accordingly expected that the dwarf goby invasion will have a strong impact on the species community and all associated ecological processes of the River Rhine and associated water bodies. Since coordinated monitoring and management of fish communities is highly benefiting from the integration of genetic and genomic analyses (Deiner et al., 2017; Pont et al., 2023; Tsuji et al., 2022), there is an urgent need for genomic resources of the dwarf goby, with a high-quality reference genome as the fundamental basis for high-resolution analyses. A reference genome is essential, serving as a detailed map of the genetic material and enabling in-depth population genomic analyses. These analyses can also help reconstruct the species’ invasion history and identify invasion routes (Jaspers et al., 2021), as well as to understand processes of rapid adaptation to local conditions in the novel environment (Szűcs et al., 2017; Yin et al., 2021).

## Material and methods

### Samples, DNA and Sequencing

Two adult individuals of the Caucasian dwarf goby *Knipowitschia* cf. *caucasica* (Figure 1) have been sampled in the River Rhine back water channel Bislich-Vahnum (North-Rhine Westphalia, Germany) in summer 2021 (sampling permission 602/00038/21 from the ULB Kreis Wesel 25.03.2021). Due to their morphological distinctness, morphological identification was straightforward. Fish were narcotised with Tricaine Methanesulfonate (MS-222) and then transferred to liquid nitrogen and preserved at -80°C.

**Figure 1.**
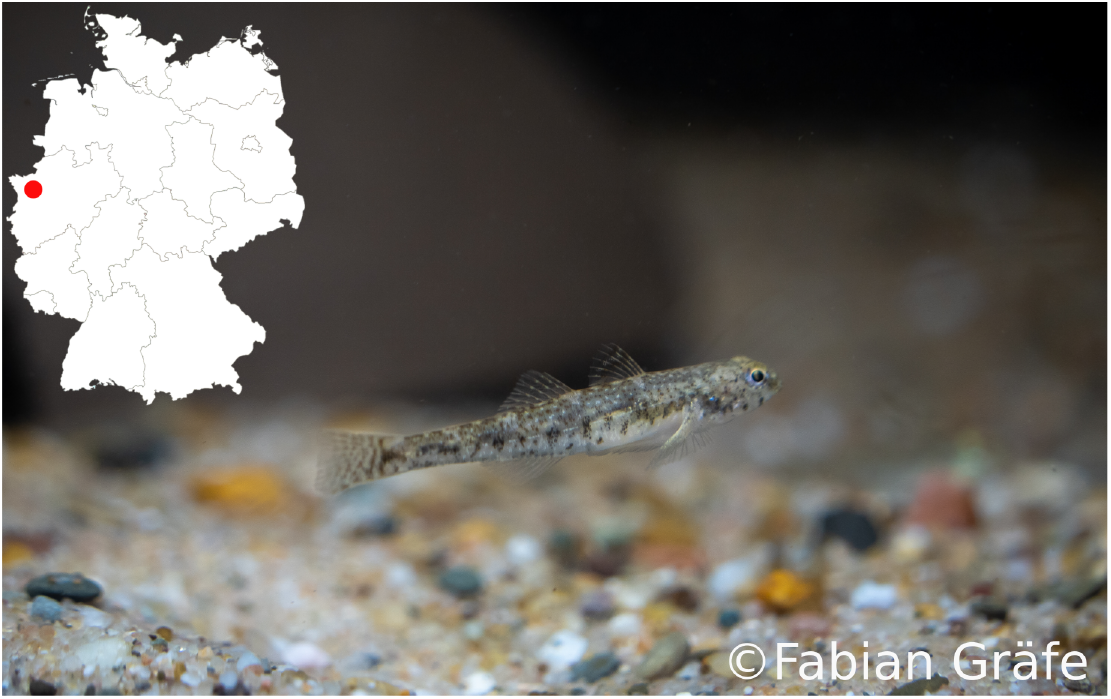
Photo of the sequenced goby species *Knipowitschia* cf. *caucasica* and sampling location (red dot) in Germany. Photo taken by Fabian Gräfe.

The genome of *Knipowitschia* cf. *caucasica* was sequenced by using a combination of PacBio Sequel IIe sequencing in CLR mode (Genome Technology Center (RGTC) at Radboudumc, Nijmegen, The Netherlands), Illumina NovaSeq 6000 sequencing with paired-end 150 bp (PE150) and Omni-C sequencing (Dovetails Ge-nomics). High-molecular-weight (HMW) DNA used for PacBio and Illumina NovaSeq sequencing was extracted from muscle tissue of one single individual following the phenol/chloroform extraction protocol (Sambrook and Russel, 2001). The proximity ligation library was constructed using the Dovetail® Omni-C® Kit (Dovetails Genomics) with a second individual (124.4 mg of muscle tissue as input material) and processed according to the Omni-C Proximity Ligation Assay protocol version 1.0. The concentration and purity of the DNA was assessed using a Nanodrop spectrophotometer and Qubit Fluorometer with the Qubit dsDNA High Sensitivity Assay kit. Fragment size distribution was evaluated by running the DNA sample on the Tapestation 2200 system to ensure that most DNA molecules were larger than 30 kb.

RNA was extracted from different tissues (gills, gonads, skin, liver, eyes, muscle) from one male individual by using the Quick-RNA MiniPrep plus Kit (Zymo Research, USA). Tissues were placed in tubes filled with 800 µl DNA/RNA shield and lysed via bead beating (speed 4 M/S, 1×30s and 1×10s). RNA extraction was done accordingly to the manufacturers protocol, with elution in 70 µl. The concentration and purity of the RNA was primarily assessed using a Nanodrop spectrophotometer and samples were pooled in the same concentration for a final sample. Quality of the sample was checked with Agilent 5400 bioanalyzer. A mRNA library with poly A enrichment was prepared and sequenced on a Illumina Novaseq machine to yield 50 millions pairs of 150-bp reads.

### Genome assembly and annotation

Illumina read quality was visualized with FastQC v0.11.9 (Andrews, 2010). Raw Illumina reads were trimmed for quality and adpaters were removed using Trimmomatic v0.39 (Bolger et al., 2014) with options LEADING:3 TRAILING:3 SLIDINGWINDOW:4:20 MINLEN:50. PacBio long reads were converted into fastq files with samtools 1.13 (Danecek et al., 2021) and assembled with three genome assembly tools including Flye 2.9 with default parameter settings (Kolmogorov et al., 2019), wtdbg2 with -L set to 5000 recommended for PacBio reads (Ruan and Li, 2020) and Raven v1.7.0 with default parameters (Vaser and Šikić, 2021). All three assemblies were compared with SeqKit stats (Shen et al., 2016) and assembly-stats (Challis, 2017). Assembly size, contiguity, BUSCO completeness and *k*-mer completeness were further checked for the two best genome assemblies (flye and raven). Ortholog completeness was analysed by using BUSCO v5.5.2 (Manni et al., 2021) with actinopterygii_odb10 (Creation date: 2021-02-19, number of genomes: 26, number of BUSCOs: 3640). *K*-mer completeness was analysed by using the tool KAT (Mapleson et al., 2016) and the KAT comp module. The best assembly with regards to assembly size, BUSCO completeness and *k*-mer completeness was used for further downstream anaylses.

To obtain chromosome resolution of the assembly, we used Omni-C reads for scaffolding of the contigs using instaGRAAL with parameters -l 5 -n 100 -c 1 -N 5 (Baudry et al., 2020). Automatic curation was done with instagraal-polish and using the -j parameter to indicate the number of N’s to put in the gaps (-j NNNNNNNNNN). For further downstream analyses the instaGRAAL assembly file was filtered for contigs larger than 10 Mb with BBmap (BBtools 2013) to only use chromosome-level scaffolds. Gaps created during the scaffolding process were closed with PacBio data using TGS-GapCloser (Xu et al., 2020) with –tgstype pb without error correction (–ne). Gap-filled scaffolds were polished with HyPo (Kundu et al., 2019) by using mapped short (Illumina) and long reads (PacBio). The genome was analyzed using BlobToolKit 4.0.7 including BUSCO scores (Challis et al., 2020). A Omni-C map for the final assembly was produced using hicstuff (Matthey-Doret et al., 2020) for contact map. To assess the assembly metrics, the estimated assembly completeness was calculated with KAT (Mapleson et al., 2016). BUSCO completeness of the final genome was analysed by using BUSCO v5.4.7 (Manni et al., 2021).

Repetitive elements were identified *de novo* with RepeatModeler version 2.0.1. Repetitive DNA and soft-masking was performed with RepeatMasker version 4.1.1 (Smit et al., 2013) using the repeat library previously identified via RepeatModeler and skipping the bacterial insertion element check (-no_is) and run with rmblastn version 2.10.0+ (Flynn et al., 2020). The masked genome assembly was used for structural annotation together with the Illumina short reads and a protein database using the BRAKER3 pipeline (Gabriel et al., 2023). The protein database consisted of the Metazoa subset from the partitioning of the OrthoDB v.11 (Kuznetsov et al., 2022) combined with protein data from three closely related species including *Mugilogobius chulae* (GCA_038363315.1), *Boleophthalmus pectinirostris* (GCF_026225935.1) and *Periophthalmus magnuspinnatus* (GCF_009829125.3) downloaded from NCBI.

For the functional annotation of the final genome assembly with InterProScan-5.59-91.0 (Jones et al., 2014) the needed sequencing data were uploaded to the Galaxy web platform provided by the Galaxy Community (2022), and we used the public server at usegalaxy.eu. The following parameters were set for the InterProScan run: -dp –seqtype p –applications TIGRFAM, FunFam, SFLD, SUPERFAMILY, PANTHER, Gene3D, Hamap, PrositeProfiles, Coils, SMART, CDD, PRINTS, PIRSR, PrositePatterns, AntiFam, Pfam, MobiDBLite, PIRSF –pathways –goterms.

Moreover, we extracted the mitochondrial genome from the assembly by using MitoHiFi 3.2 (Uliano-Silva et al., 2023) with MitoFinder (Allio et al., 2020).

To test the quality of genome scaffolding, chromosome-scale collinearity analysis was conducted between the here presented *Knipowitschia* cf. *caucasica* (2n=46) and the yellowstripe goby *Mugilogobius chulae* (2n=44) by using MCScanX (Y Wang, Tang, DeBarry, et al., 2012). Belonging to the family of Gobiidae, both species have chromosome level genomes with available annotations and genomes of similar size with 957 Mb of *Knipowitschia* cf. *caucasica* (this study) and 1 Gb *M*.*chulae*. Genome data for *M. chulae* were downloaded from NCBI (GCA_038363315.1). The generation of .gff input files was done by using the mkGFF3.pl program in the MCScanX_protocol (Y Wang, Tang, X Wang, et al., 2024) with the genome.gff files and CDS_from_genomic.fna files. A protein database was created from both species by using their protein fasta files with BlastP and makeblastdb (-dbtype prot). An all-against-all BLASTP run was conducted with an E-value cutoff of 1e-10 and keeping the best five non-self-hits (–num_alignments 5). Gff files and blast output files of both species were concatenated and a collinearity analysis was conducted by by using MCScanX, which creates a .collinearity file (containing pairwise colinear blocks) and .tandem file (listing all consecutive repeats). Synteny was visualized by using the MCScanX downstream tools (bar_plotter.java, dot_plotter.java) creating bar plots and dot plots. A dual synteny plot was created with SynVisio (Bandi and Gutwin, 2020).

## Results

The long-read assembler Flye (Kolmogorov et al., 2019) generated the best out of three obtained assemblies with regards to the assembly size, contiguity, BUSCO completeness (Table 1) and k-mer completeness. This assembly consisted of 16,269 contigs (966 Mb total contig length). Contigs displayed high contiguity with an N50 length of 149.4 kb. The integrity of the flye assembly was demonstrated by 90.52% BUSCO gene completeness (single 89.09%, duplicated 1.43%) using the actinopterygii_odb10 reference set. Thus, we used the flye assembly for further downstream analyses to generate the final complete genome of *Knipowitschia* cf. *caucasica*.

**Table 1.** Genome assembly data for three tested assembly tools (flye, raven, wtdbg2) including info on assembly size, contiguity and BUSCO completeness (based on actinopterygii_odb10). BUSCO completeness was not tested for wtdbg2 assembly due to low contig N50 length. NA: not available

|  | flye | raven | wtdbg2 |
| --- | --- | --- | --- |
| span (bp) | 996,499,675 | 985,684,880 | 787,966,447 |
| N (%) | 0 | 0 | 0 |
| GC (%) | 42.90 | 43.16 | 42.59 |
| AT (%) | 57.10 | 56.84 | 57.14 |
| contig count | 16,269 | 14,297 | 20,132 |
| longest contig (bp) | 1,996,640 | 742,234 | 456,979 |
| contig N50 length (bp) | 149,364 | 92,290 | 66,168 |
| contig N50 count | 1,764 | 2,996 | 3,556 |
| contig N90 length (bp) | 27,499 | 32,547 | 17,751 |
| contig N90 count | 7,557 | 9,903 | 12,326 |
| BUSCO completeness (%) | 90.52 | 84.09 | NA |
| BUSCO single (%) | 89.09 | 82.23 | NA |
| BUSCO duplicated (%) | 1.43 | 1.87 | NA |

Contigs obtained by the Flye assembler were scaffolded into 23 chromosome-scale scaffolds (956.6 Mb in length, 98.87% assembly length, 43 Mb scaffold N50 length) using the Omni-C data (Figure 2B). BUSCO scores (Manni et al., 2021) of the final genome assembly after gap-closing and polishing resulted in 92.3% gene completeness (single 91.1%, duplicated 1.2% using the actinopterygii_odb10 reference set (Figure 2A).

**Figure 2.**
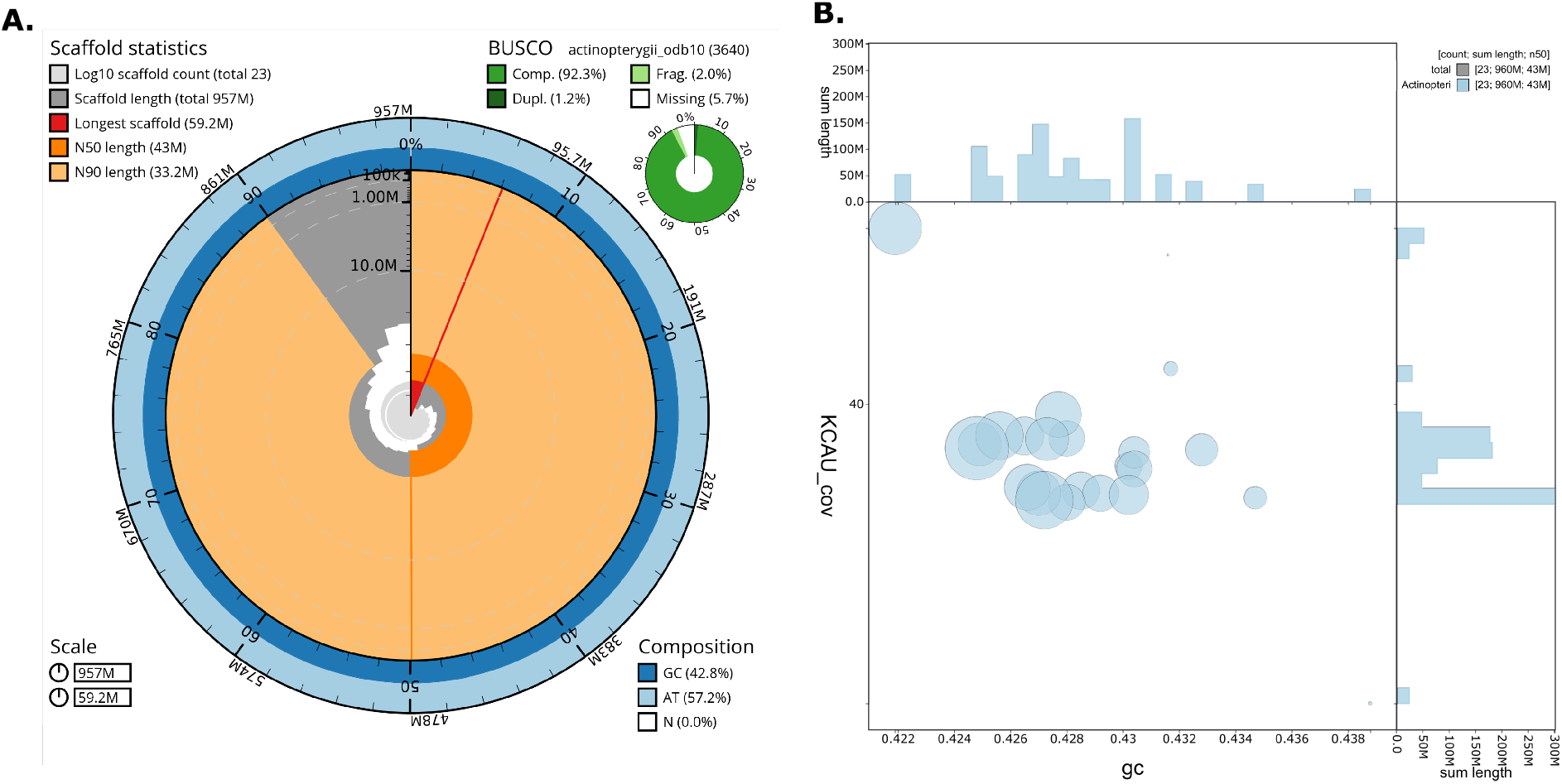
**A**. Snailplot produced by Blobtoolkit2 showing assembly metrics, including scaffold statistics (top left), BUSCO completeness based on the actinopterygii_odb10 reference set (top right), and base composition (bottom right) shown by dark/light blue rings. The inner radial axis (gray) shows the length of each contig in descending order. Dark orange and light orange portions represent the N50 and N90 scaffold lengths, respectively. The genome has a total length of 957 Mb with a maximum contig length of 59.2 Mb (shown in red). Distribution of scaffold lengths is shown in dark grey with the plot radius scaled to the longest scaffold present in the assembly (59,227,272 bp, shown in red). Orange and pale-orange arcs show the N50 and N90 scaffold lengths (43,016,011 bp and 33,193,346 bp), respectively. **B**. BlobToolKit GC-coverage plot. Scaffolds are color-coded based on their phylum, while circles are scaled proportionally to scaffold length. Histograms depict the distribution of the total scaffold length along each axis.

Gene annotation predicted a total of 26,959 genes of which 555 genes were duplicated. The final genome revealed a total of 26,404 transcripts distributed across 21,443 loci, with 23,210 transcripts characterized as multi-exon, of which 26,260 transcripts were functionally annotated. BUSCO analysis recovered 89.5% BUSCO gene completeness (single 73.5%, duplicated 16.0%) using the actinopterygii_odb10 reference set. KAT analysis based on Illumina reads and the collapsed assembly showed two peaks of *k*-mer multiplicity; a heterozygous peak at 24X and a homozygous peak at 48X, with almost all *k*-mers represented exactly once in the homozygous peak of the assembly as expected (Figure 3A). The estimated assembly completeness of the final assembly calculated with KAT was 96.64%. Chromosome-scale scaffolds were labeled by decreasing size. The remaining 1.03% unplaced sequences were smaller than 10,000 kb. The chromosome-level scaffolds showed relatively consistent contact patterns, representing well individualized entities in the contact map (Figure 3B). The mitochondrial genome has also been assembled and is 16,377 bp in length.

**Figure 3.**
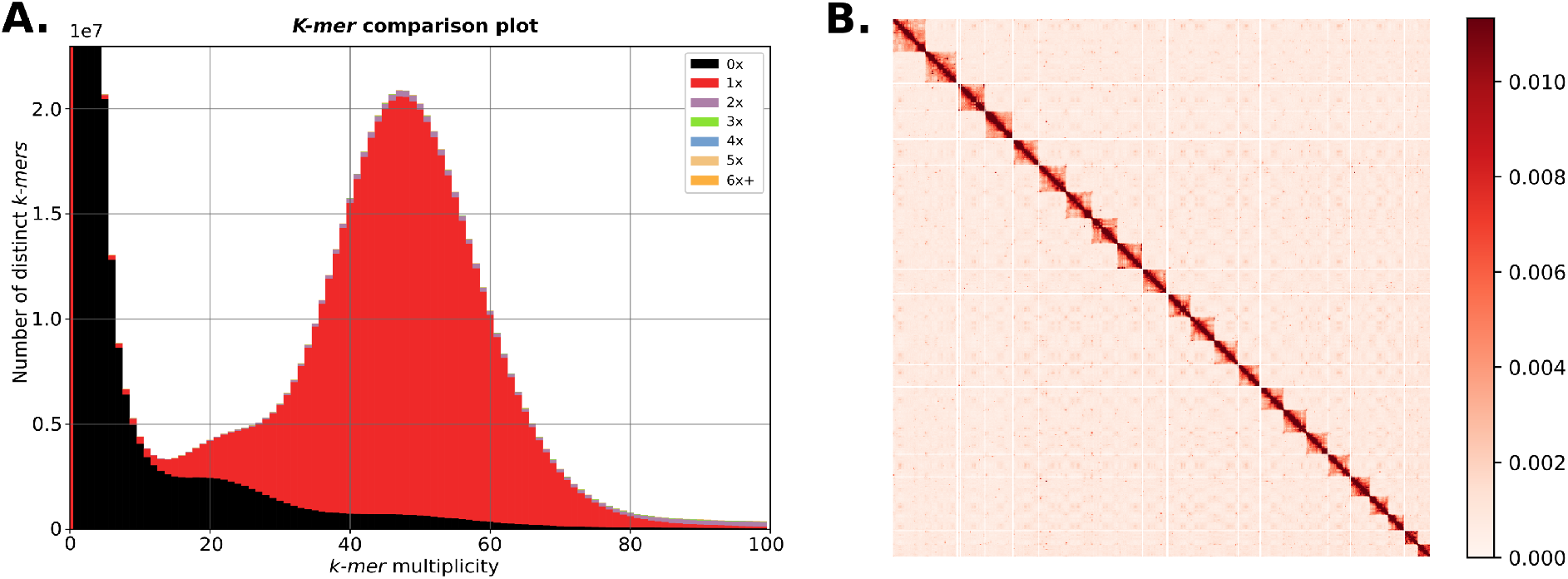
**A**. KAT comparison of the *k*-mers in the Illumina dataset versus the final genome assembly. **B**. Omni-C contact map, with a binning of 3000 and normalization, for the final genome assembly. All 23 chromosomes are shown in order of size from left to right and top to bottom.

The repeat content was identified via RepeatModeler and resulted in a non-redundant library of 2,812 consensus sequences of repeat families (Supplement https://doi.org/10.5281/zenodo.10784873). These repeats accounted in total for 53.08 % (507.8 Mb) of the assembled genome.

Results of chromosome-scale collinearity analysis between *K. caucasica* and *M. chulae* showed high synteny conservation (37.07% of shared colinear genes), suggesting a high quality of genome scaffolding for *Knipowitschia* cf. *caucasica* (Figure 4A-C).

**Figure 4.**
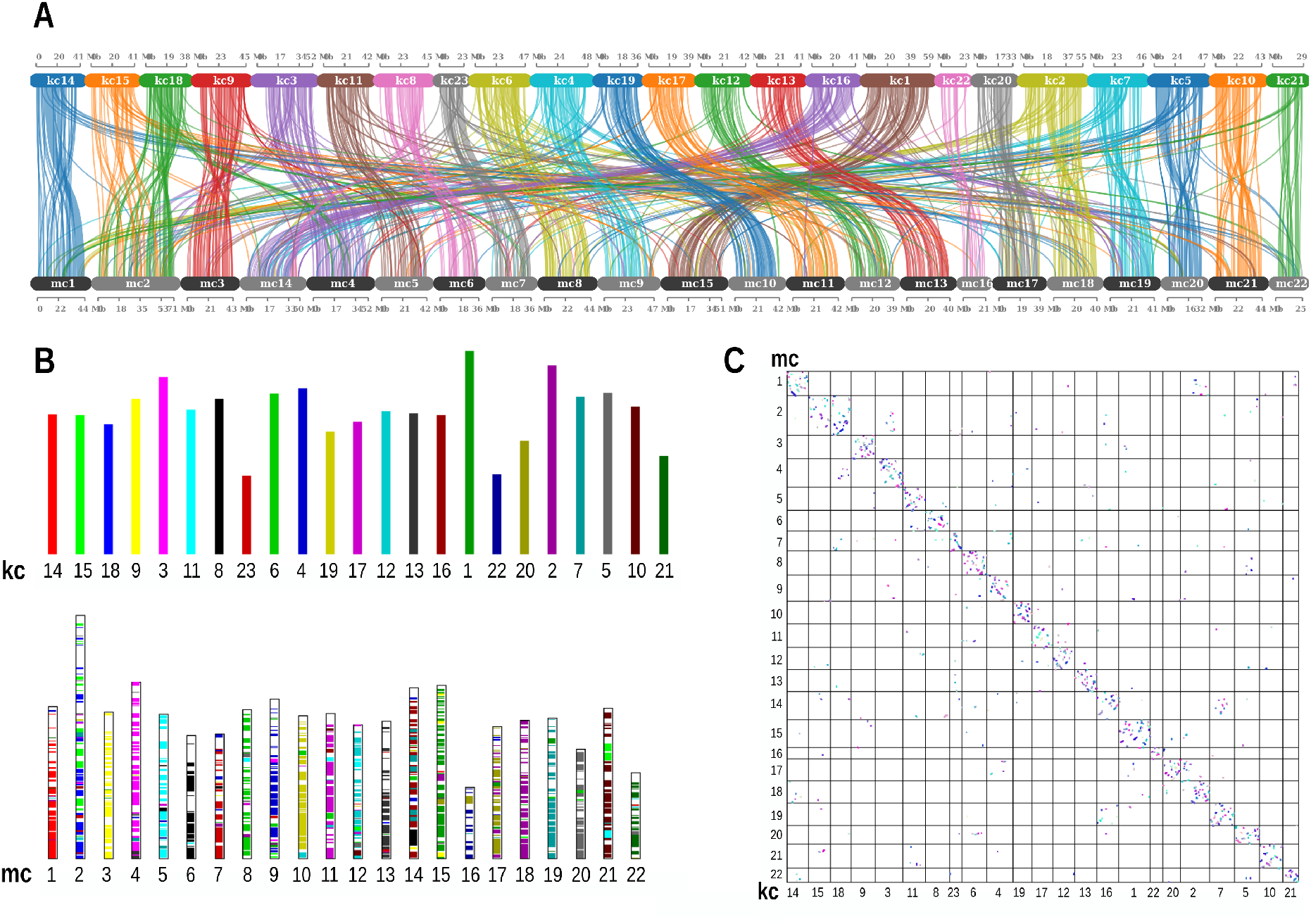
**A**. Dual synteny plot of chromosome sets from *Knipowitschia* cf. *caucasica* (kc) and *Mugilogobius chulae* (mc). Lines connect pairs of colinear genes between the two sets of chromosomes. **B**. Synteny visualization with bar plots of *Knipowitschia* cf. *caucasica* (kc) chromosomes (upper panel, each chromosome represented by unique color) and *Mugilogobius chulae* (mc) chromosomes (lower panel). The mc set of chromosomes contains colinear blocks denoted by the colors of the chromosomes where their paired colinear blocks are located. **C**. Dot plot of *Knipowitschia* cf. *caucasica* (x-axis) and *M. chulae* (y-axis) chromosomes. Dots represent pairs of colinear genes between the two sets of chromosomes.

## Discussion

With the benefit of PacBio sequencing, genome assembly was performed with three different approaches, and we selected the most performant software (Flye) for contig construction. Here we present the complete genome sequence of the invasive Gobiidae species *Knipowitschia* cf. *caucasica*, generated using Illumina and PacBio platforms, to achieve an assembly of approximately 956.58 Mb (scaffold N50 of 43 Mb) and high contiguity with 26,404 transcripts of which 26,260 transcripts were functionally annotated. In comparison, the benthic round goby *Neogobius melanostomus* genome is 1 Gb in size with a gene annotation prediction of 38,773 genes and 39,166 proteins reaching a BUSCO completeness of 86.9 % (Actinopterygii) (Adrian-Kalchhauser et al., 2020), while for the blue-spotted mudskipper *Boleophthalmus pectinirostris* the genome size is 957.8 Mb with 22,685 genes (Bian et al., 2024).

Studying population genomics and establishing reference genomes for invasive species plays a crucial role in advancing our comprehension of biological invasions and aids in the proactive detection and management of these invasive species. Despite a global increase in invasion rates, the field of “invasion genomics” is still in its infancy. In a comprehensive review, it was noted that only 32% of species listed on the International Union for Conservation of Nature’s “100 Worst Invasive Alien Species” have undergone studies utilizing population genomic data. Furthermore, for over 50% of the species on this list, a reference genome is yet to be established (Matheson and McGaughran, 2022). Therefore, the assembly of reference genomes for invasive species is imperative and essential to unravel the role of genome-driven processes in facilitating invasion, and making them publicly available serves as a crucial step.

## Acknowledgements

We thank the Genome Technology Center (RGTC) at Radboudumc for the use of the Sequencing Core Facility (Nijmegen, The Netherlands), which provided the PacBio SMRT sequencing service on the Sequel IIe platform. We acknowledge the Regional Computing Center of the University of Cologne (RRZK) for providing computing time on the DFG-funded [Funding number INST 216/512/1FUGG] High Performance Computing (HPC) system CHEOPS as well as support. We also thank the ULB Kreis Wesel for granting us permission to sample the fish at Bislich-Vahnum, North-Rhine-Westphalia, Germany.

## Fundings

AMW acknowledges funding of her junior professorship as part of the BMBF funded “Bund-Länder-Programm This project is also part of the Biodiversity Genomics Center Cologne (BioC^2^) funded by the Excellence Research Support Programm of the University of Cologne (UoC Forum).

### Data, script, code, and supplementary information availability

The genome sequence of *Knipowitschia* cf. *caucasica* (caucasian dwarf goby) is released openly for reuse. The *Knipowitschia* cf. *caucasica* genome sequencing is part of the LTER project REES (https://deims.org/554de3a9-1ad9-46e9-9b70-f6e25a799876). All raw sequence data and the assembly have been deposited at the European Nucleotide Archive (ENA) under Project accession number PRJEB58922: https://identifiers.org/ena.embl/PRJEB58922. The repeat library is depsoited at Zenodo (https://doi.org/10.5281/zenodo.10784873)

